# The Transcriptional Architecture of Bacterial Biosynthetic Gene Clusters

**DOI:** 10.1101/2025.03.18.644061

**Authors:** Silvia Ribeiro Monteiro, Yasmine Kerdel, Julianne Gathot, Sébastien Rigali

## Abstract

Bacteria produce diverse bioactive metabolites with ecological and pharmaceutical importance. These compounds are synthesized by biosynthetic gene clusters (BGCs), whose expression is tightly regulated. While many studies have examined the factors influencing BGC expression, including transcription factors (TFs) and environmental signals, the regulatory architecture governing BGCs expression remains largely unexplored. In this meta-analysis, we collected experimental datasets of bacterial transcription factor binding sites (TFBSs) to unveil i) the functional gene categories preferentially targeted by TFs, ii) the regulatory coverage based on cluster organization, iii) the positional distribution of TFBSs, and iv) the binding strength of TFs. Our analysis reveals a regulatory strategy where global TFs primarily target pathway-specific TFs when present, aligning with a “one-for-all” strategy ensuring cluster-wide expression control. Additionally, examination of the organization of TFBS-associated genes identified distinct transcriptional strategies: regulatory genes are frequently monocistronic, while biosynthetic genes tend to be co-transcribed in operons to guarantee biosynthesis efficiency. The positional distribution of TFBSs highlights a strong enrichment in the upstream regions of genes optimizing their role in gene regulation. Finally, assessment of TF-TFBS interaction strength suggests that TFBSs within BGCs exhibit lower binding affinities compared to those associated with core regulon genes that reside outside BGCs, allowing greater regulatory flexibility in response to multiple environmental cues. These findings provide new insights into the regulatory principles shaping BGC expression and would help predict conditions for activating cryptic BGCs, facilitating the discovery of novel bioactive compounds through targeted culture and engineering strategies.

## Introduction

Bacteria are remarkable producers of specialized metabolites (polyketides, nonribosomal peptides, terpenoids, aminoglycosides, quinones…) which are small organic compounds critical for survival in natural habitats. These compounds display an extraordinary structural and bioactive diversity (e.g. antimicrobials, toxins, anti-oxidants, metal-chelators, signaling molecules…) which have provided useful drugs or leads for treating infections and cancer chemotherapy amongst other applications ^1,2^. Only an estimated 3% of the natural products encoded in bacterial genomes have been experimentally characterized, highlighting the immense potential for discovering numerous valuable compounds in the future ^3^. Their production depends on biosynthetic gene clusters (BGCs), which are groups of genes that work together to coordinate biosynthesis, transport, regulation, and, when mandatory, self-resistance mechanisms to the compound they produce. The Minimum Information about a Biosynthetic Gene cluster (MIBiG) provides a standardized framework for annotating gene groups, enabling the systematic classification of genes according to their role (Figure 1A) ^4^. The genes for biosynthesis in BGCs can be of two types, either “core” or “additional”, each playing distinct roles in metabolite production. Core biosynthetic genes encode the enzymes directly responsible for constructing the backbone of the metabolite which can then be chemically modified by enzymes encoded by additional biosynthetic genes. In some cases, additional biosynthetic genes also encode enzymes involved in metabolic pathways that generate precursors and building blocks used by core biosynthetic enzymes in order to ensure the timely supply of substrates to construct the metabolite’s primary structure ^5,6^. The presence of genes for building block biosynthesis can also help predicting structural features of compounds associated with cryptic BGCs thereby facilitating the discovery of novel natural products ^7^. Coordinated expression in BGCs also includes genes encoding transport-related proteins with diverse functions such as i) secreting the natural product to fulfill its role in the surrounding environment, ii) importing the metabolite for intracellular processes, iii) providing a resistance mechanism when the metabolite is toxic to the producer ^8,9^, iv) importing building blocks ^10^, and v) importing signaling molecules to trigger or repress the expression of the BGC.

**Figure 1.**
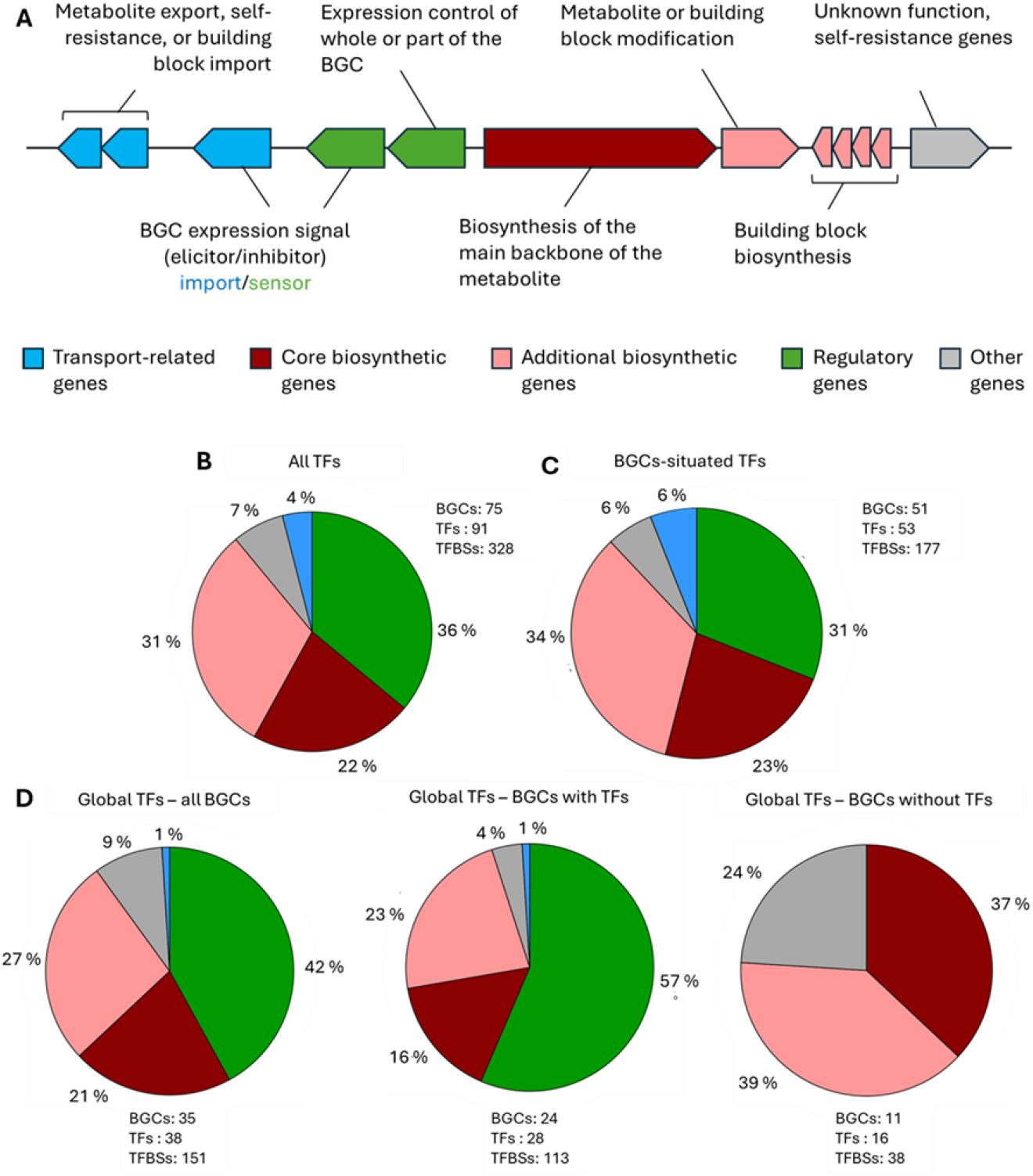
Occurrence of TFBS according to the gene functional categories in BGCs. **A**. Functional categories and the roles of genes in BGCs. The gene functions are color-coded according to the scheme applied across MIBiG entries. **B**. Percentage of gene functional categories targeted by both global and BGCs-situated TFs in all BGCs. **C**. Percentage of gene functional categories targeted by BGCs-situated TFs. **D**. Percentage of gene functional categories targeted global TFs for all BGCs (left panel), for BGCs that include pathway-specific TFs (center panel), and for BGCs that do not include specific TFs (right panel).

The interplay between biosynthetic and transport-related genes illustrates how bacteria tightly regulate their secondary metabolism. By definition, *specialized* metabolites are produced in response to specific environmental cues and/or at a precise stage of bacterial growth or developmental program to meet particular needs ^11,12^. Consequently, the expression of BGCs is subjected to intricate multilevel transcriptional ^13–15^ and post-transcriptional controls ^16^, reflecting both the diverse ecological niches in which microorganisms evolve and the evolutionary pressures they face. Understanding the regulation of BGCs is essential for multiple reasons. Since BGC-associated compounds often contribute to microbial competition and survival, identifying the signals and molecular mechanisms that control their expression provides insight into how bacteria interact with their environment. To ensure proper spatiotemporal production of bioactive natural products, BGCs frequently include genes encoding transcription factors (TFs) that regulate the expression of the BGC they reside in, referred to as cluster-situated and/or pathway-specific regulators ^17^. In addition to these specialized TFs, BGCs’ expression is also directly influenced by pleiotropic and global TFs, which exert broader control over multiple biological processes ^13– 15,18,19^. Global regulators often act as molecular bridges, linking environmental cues, such as sensing host-related signals, to the production of metabolites, including those involved in host colonization and virulence ^20^. From an applied perspective, many bioactive compounds are produced inefficiently in their native hosts. Modulating BGC regulation can enhance yields, making large-scale production more cost-effective. Additionally, many bacteria harbor silent or weakly expressed BGCs that remain inactive under standard laboratory conditions. Understanding their regulation allows researchers to activate these clusters, unlocking novel secondary metabolites with potential pharmaceutical applications^21,22^. With more than 2,000 bacterial BGCs experimentally characterized ^23^, substantial data now exist to deepen our understanding of how BGC expression is controlled. Key questions remain about the functional gene categories targeted by global TFs within a BGC, the positional distribution of transcription factor binding sites (TFBSs) across the different types of genomic regions, and the interaction strength between TFs and their binding sites. In this meta-analysis, we collected experimental datasets of hundreds of TFBSs to shed light on the general and specific features of the regulatory architecture governing the expression control of bacterial BGCs.

### Functional targeting: Genes’ function in BGC targeted by transcription factors

Coordinated expression of all gene types within a BGC is rational for several reasons. First, overexpression of core biosynthetic enzymes alone is ineffective if the uptake or synthesis of their substrates by transporter or additional biosynthetic genes is not simultaneously enhanced. Second, failing to concomitantly enhance the expression of export-related genes could lead to intracellular metabolite accumulation, potentially overwhelming the organism’s self-resistance mechanisms. But how do global TFs coordinate the expression of all genes within BGCs? Do they evenly distribute TFBSs across all transcriptional units, or do they primarily target the BGC-situated TF, which typically governs the entire cluster’s expression? To address this, we analyzed experimental datasets of TFBSs identified within BGCs using techniques that provide direct evidence of TF-TFBS interactions, such as DNAse footprinting assays, ElectroMobility Shift Assays (EMSA), Chromatin ImmunoPrecipitation on Chip (ChIP-on-chip), or/and Chromatin ImmunoPrecipitation sequencing (ChIP-seq) assays. The results of a literature survey on 91 TFs (38 global and 53 pathway-specific TFs) associated with the control of 75 BGCs across 17 bacterial genera, allowed us to collect 328 TFBSs. The distribution of TFBSs according to the different functional categories in BGCs is presented in Figure 1B. Overall, the binding sites of TFs are most frequently associated with the genes of the regulatory functional category (36%), followed by additional biosynthetic genes (31%), core biosynthetic genes (22%), genes with “other or unknown” functions (7%), and finally genes for transport-related proteins (4%) (Figure 1B). When analyzing only the TFBSs of cluster-situated TFs, the distribution of the binding sites is very similar with 34%, 31%, and 23% of TFBSs being associated with additional biosynthetic genes, regulatory genes, and core biosynthetic genes, respectively (Figure 1C). Among the 31% of TFBSs linked to TFs, 38% were located near their corresponding TF, suggesting autoregulation of their own expression via positive or negative feedback loops, depending on whether they function as activators or repressors. The targeting of genes encoding other regulatory proteins for 62% of TFBSs confirms the existence of regulatory cascades, where several TFs within a BGC occupy distinct hierarchical roles to coordinate the expression control of the entire cluster ^19^. Interestingly, when considering only global TFs, the proportion of TFBSs targeting the regulatory category increases to 42% (Figure 1D, left panel). However, not all BGCs contain a dedicated pathway-specific TF. When we reanalyzed only BGCs that include a regulatory protein, the bias toward targeting regulatory genes became even more pronounced, rising to 57% (Figure 1D, center panel). This enrichment is especially striking given the relatively low proportion of regulatory genes compared to biosynthetic genes within BGCs. These findings suggest a “one-for-all” regulatory strategy: since pathway-specific regulators often activate the entire BGC ^17^, targeting the “regulation” functional category ensures global TFs to exert comprehensive transcriptional control over the whole cluster. When instead there is no regulatory proteins in a BGC, TFBSs of global TFs are almost equally associated with additional biosynthetic genes (39%) and core biosynthetic genes (37%), followed by genes with other or yet unknown functions (24%) (Figure 1D, right panel). Overall, our results highlight a preferential regulatory mechanism where global TFs primarily target pathway-specific regulators when present but shift their focus to biosynthetic genes in their absence.

### Organization of genes targeted by transcription factors

Bacterial genes involved in the same biological process are often organized into operons or co-expressed transcription units (TUs) within a cluster. Consequently, a TFBS associated with one gene can influence downstream genes transcribed in the same direction, ensuring synchronized expression. This organization enables bacteria to regulate entire biosynthetic processes in a coordinated manner, and understanding operon structures helps predict how regulatory inputs affect multiple genes simultaneously. Using existing expression data (e.g., RT-PCR), we determined the number of genes across the various functional categories that are co-transcribed with the gene where TFBSs were identified. Among the 400 genes retrieved from 69 BGCs, we identified 151 TUs, 80 (53%) being monocistronic genes and 71 (47%) organized in operon (Figure 2A). The number of genes included in an operon varied from 2 to a maximum of 19 genes (Figure 2B). Regulatory genes are the most abundant monocistronic genes (Figure 2A), as they often require precise and independent control, separate from biosynthetic operons. Monocistronic transcription allows finely tuned expression in response to environmental signals prior to affect other genes in the cluster. As shown in Figure 1D, many pathway-specific TFs autoregulate their own expression through positive or negative feedback. This mechanism is more efficient when the TF is transcribed independently, enabling a rapid response to fluctuations in metabolite levels or environmental cues. In contrast, additional biosynthetic genes belong to the functional category most commonly found within operons (Figure 2A). Many of these genes encode tailoring enzymes that introduce structural modifications essential for the final bioactive compound. Organizing them in operons alongside core biosynthetic genes ensures their coordinated expression, preventing the accumulation of incomplete intermediates and optimizing metabolic efficiency. These findings highlight distinct transcriptional strategies within BGCs: regulatory genes often function independently for precise control, while additional biosynthetic genes are co-transcribed in operons to ensure whole biosynthetic pathway progression.

**Figure 2.**
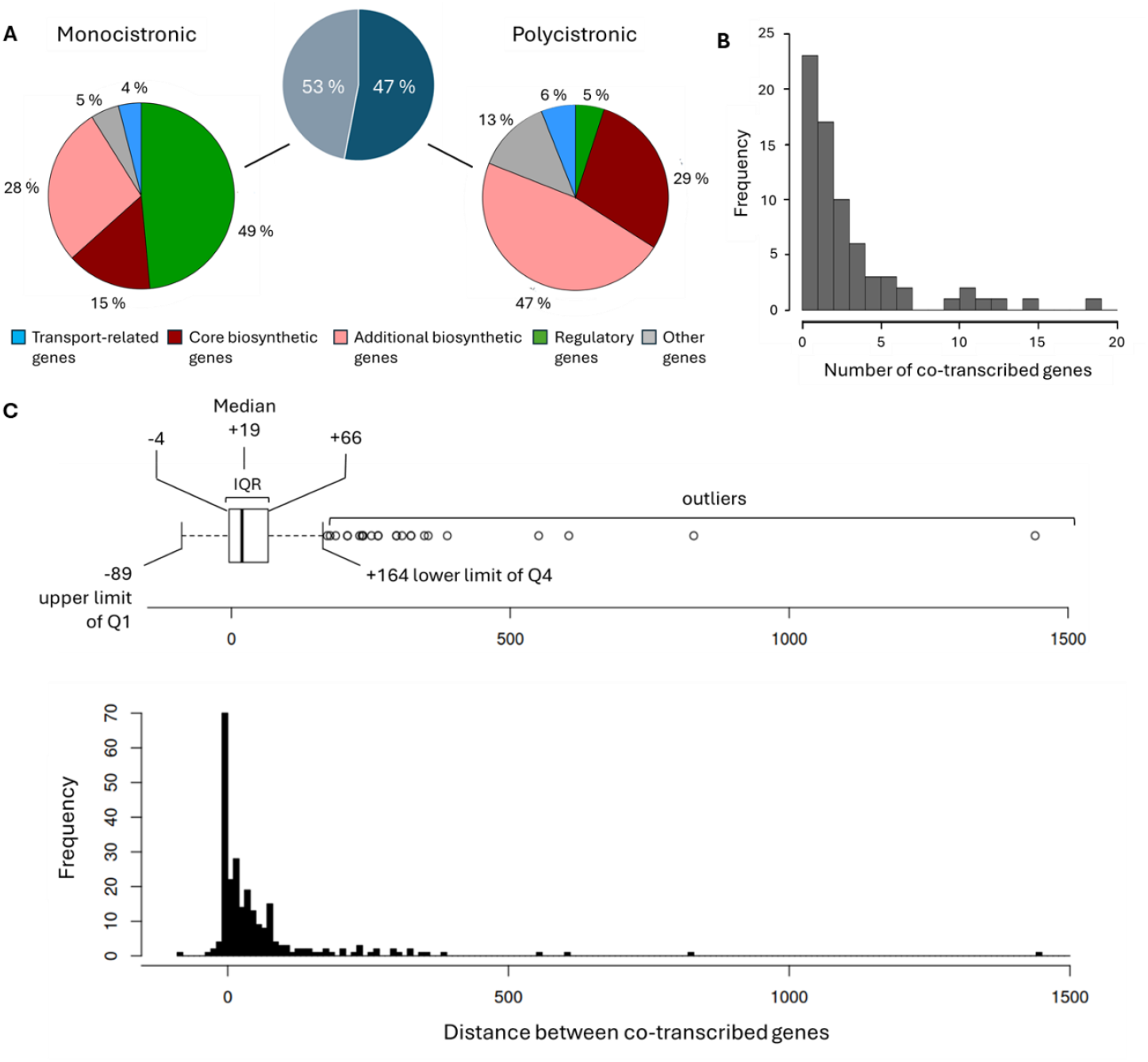
Organization of BGCs’ genes targeted by TFs. **A**. Percentage of transcription units associated with a TFBS that are either organized as monocistronic gene or in operon. **B**. Number of genes and their frequency found in BGCs’ operons. **C**. Distribution of intergenic distances between pairs of adjacent or overlapping genes in operons. (distance = Start coordinate of downstream gene - End coordinate of upstream gene +1). The edges in the boxplot indicate the 1st and 3rd quartiles (Q), and the median as center line. IQR, Inter-quartile range.

The data collected allowed us to deduce the distance tendencies of adjacent genes within operons of BGCs. The median distance between the end and start of two adjacent co-transcribed genes is 19 nucleotides (nt) with the inter-quartile range (the distance of 50% of start codons from the upstream stop codon) between nucleotide positions -4 and +66 nt relative to the translation stop codon (Figure 2C). Interestingly, there is a frequency peak at position -4, with nearly 25% of all collected distances between adjacent co-transcribed genes occurring at this specific position. The four quartile groups ranging from -89 and +164 nt as the upper and lower limits, respectively, define the distance between most (90%) co-transcribed genes in BGCs. Only 24 (10%) outlier values were found with an exceptional case of 1441 nt intergenic distance due to the insertion of an integrase gene (in opposite direction) within a TU ^24^. Based on the frequency distance distributions we provide useful information to predict the TU organization.

### Positional distribution of TFBSs in BGCs

Another key question concerns the positional distribution of TFBSs across different types of regions in a BGC, i.e. the gene coding sequences, and the intergenic regions, the latter comprising the genes’ upstream regions and the “terminator” regions (between two stop codons) (Figure 3A). Another region called “regulatory” is also occasionally stated and refers to the positions where promoters and TFBSs are likely to be found (Figure 3A). Genes’ coding and intergenic regions are easily determined based on the positions of the start and stop codons. In contrast, delineating the boundaries of the regulatory region cannot be anticipated given that the exact position of TFBSs regarding the transcriptional and the translational start sites varies widely for each TF, and according to the possible presence of one or more internal promoters within the transcriptional units they control.

**Figure 3.**
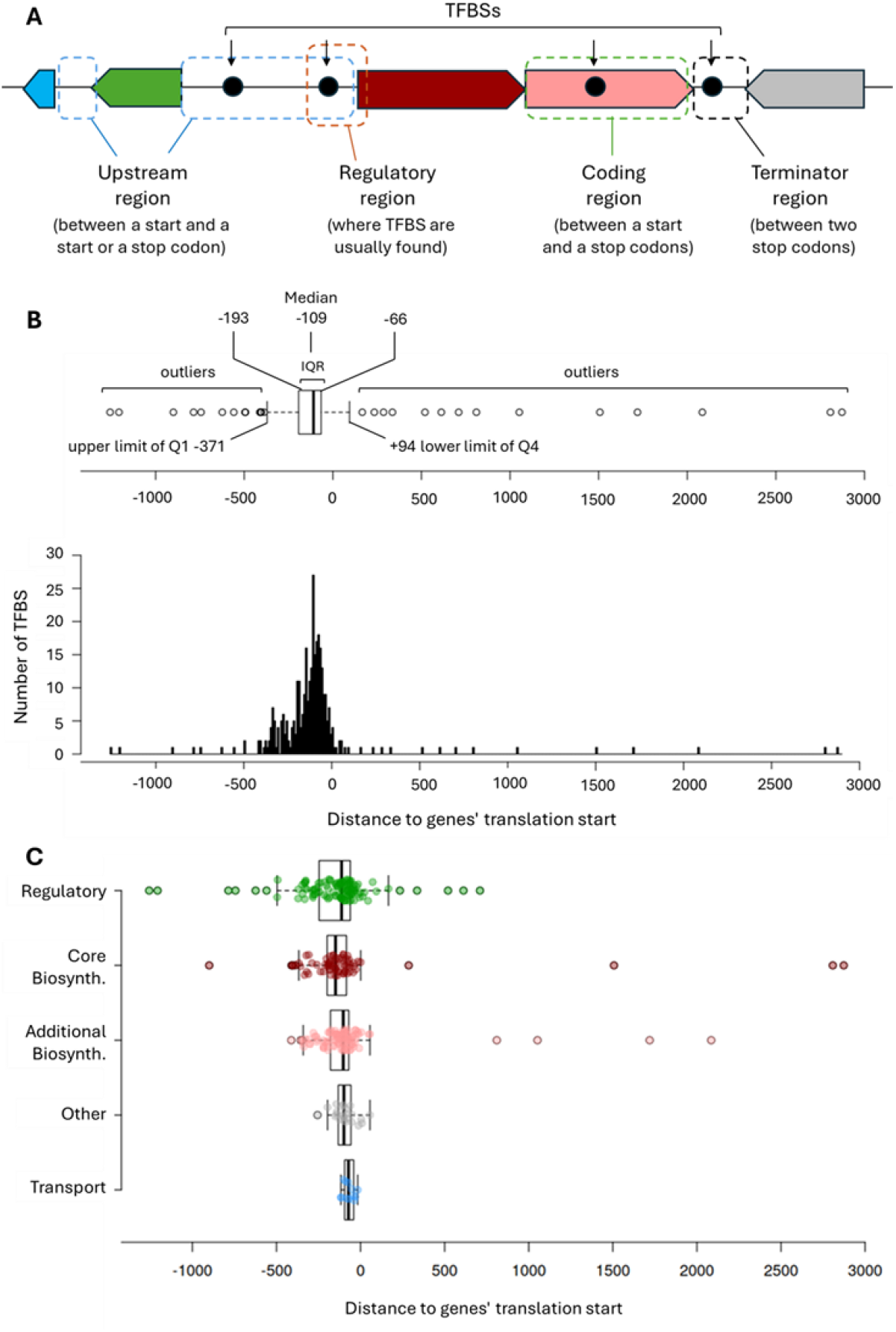
Positional distribution of TFBSs in BGCs. **A**. Definition of the different types of regions in a gene cluster. **B**. Histogram (lower panel) and Boxplot (upper panel) showing the positional distribution of all TFBSs. The edges in the boxplot indicate the 1st and 3rd quartiles (Q), and the median as center line. IQR, Inter-quartile range. C. Boxplots showing the positional distribution of TFBSs according to the genes’ functional categories.

The positional distribution of 328 experimentally validated BGC-associated TFBSs reveals a strong density in the genes’ upstream region (Figure 3B). The TFBSs distribution enabled to determine the inter-quartile range (the position of 50% of TFBSs) between nucleotide positions -192 and -66 relative to the translation start codon, with the median at position -109 nt. The four quartile groups ranging from -371 and +94 nt as the upper and lower limits, respectively, define the “regulatory” region of BGCs where most TFBSs (92%) are located. Only 14 (4%) outliers TFBSs are found upstream of and another 14 downstream of the regulatory region. These results contrast with recent genome-scale studies that have shown that model bacterial TFs exhibit similar binding patterns in both coding sequences and regulatory regions, with a widespread presence of internal cryptic promoters within coding sequences ^25^. Since BGCs often exceed 200 kb and core biosynthetic genes also being exceptionally large and organized in operons, such features may be favorable to the presence of internal promoters – and therefore internal TFBSs – in the coding sequences. However, the positional distribution of TFBSs according to the genes’ functional categories did not reveal such internal position of TFBS neither for core biosynthetic genes nor for any other functional category (Figure 3C).

### Strength assessment of the TFBS-TF interactions in BGCs

Finally, we aimed to compare the strength of interaction between a global TF and its TFBSs in BGCs versus those associated with genes outside BGCs, which are involved in broader biological processes and conserved across organisms. The rationale behind this comparison is that TFBSs of core regulon genes typically exhibit nucleotide sequences optimized for specific and tight recognition. In contrast, in BGCs whose presence is often limited to a series of species within a bacterial genus, TFBSs may be less tightly bound by the TF. This reduced affinity could stem from the need for these genes to respond to multiple environmental signals rather than their expression being exclusively regulated by a single TF, leading to deviations from the canonical TFBS sequences and therefore reducing binding affinity. To assess interaction strength, we used statistical models of TFBS preferences, applying position weight matrices to score sequence similarity to the consensus sequence following the scoring methodology described in ^26^ and calculated via the PREDetector software ^27^. These scores, normalized between 0 and 1 (with 1 representing the consensus sequence), were used to compare the strength of TFBSs within BGCs and those associated with non-BGC genes.

Overall, TFBSs within BGCs exhibit lower strength scores compared to those outside BGCs, with median scores of 0.53 and 0.84, respectively (Figure 4A). However, strength score distributions vary depending on the biological process regulated by each analyzed TF. TFs principally involved in primary metabolism, such as CebR (cellulose utilization) ^28,29^, BxlR (xylan utilization) ^30^, and DasR (chitin metabolism)^31–34^, show higher strength scores for TFBSs of core regulon genes involved in carbon source utilization than for those within BGCs. For instance, the CebR TFBSs found upstream of genes of the cellulolytic system display the perfect palindromic 14 nt sequence TGGGAGCGCTCCCA ^20^. In contrast, CebR TFBSs associated with the thaxtomin-production BGC in plant-pathogenic species consistently contain at least one mutation ^35^. These mutations weaken CebR binding, leading to a lower expression fold-change of thaxtomin biosynthetic genes upon sensing the cellotriose elicitor, compared to the stronger response of primary metabolism genes involved in cellulose byproduct import and catabolism^36^. The contrast in strength score distribution is even more pronounced for the chitin utilization regulator DasR. While TFBSs upstream of chitinase and N-acetylglucosamine utilization genes accept a maximum of three mismatches, those within BGCs frequently contain up to six mismatches, further reducing DasR binding (Figure 4B). A similar contrast in strength scores is observed for the redox-sensitive transcriptional regulator SoxR ^37^, and for TFs involved in micronutrient utilization, such as PhoP^38^, GlnR^39,40^, and Zur^41–43^, which regulate phosphate, nitrogen, and zinc sources uptake/utilization, respectively (Figure 4B). Instead, TFs typically involved in morphological and/or physiological differentiations in *Streptomyces* species such as AdpA^44,45^, AfsQ1^46^, Crp^47,48^, and MtrA^49,50^, exhibit a more homogeneous distribution of interaction strength scores, consistent with their role in specialized metabolite production control (Figure 4B). These results support our hypothesis that core regulon genes require strict regulation, relying on high-affinity TFBSs and transcriptional activation only in response to their specific substrate. In contrast, BGC-associated genes, which are occasionally regulated by these TFs, utilize sub-optimal TFBSs. This reduced binding affinity enables a more flexible expression pattern, allowing adaptation to multiple environmental signals.

**Figure 4.**
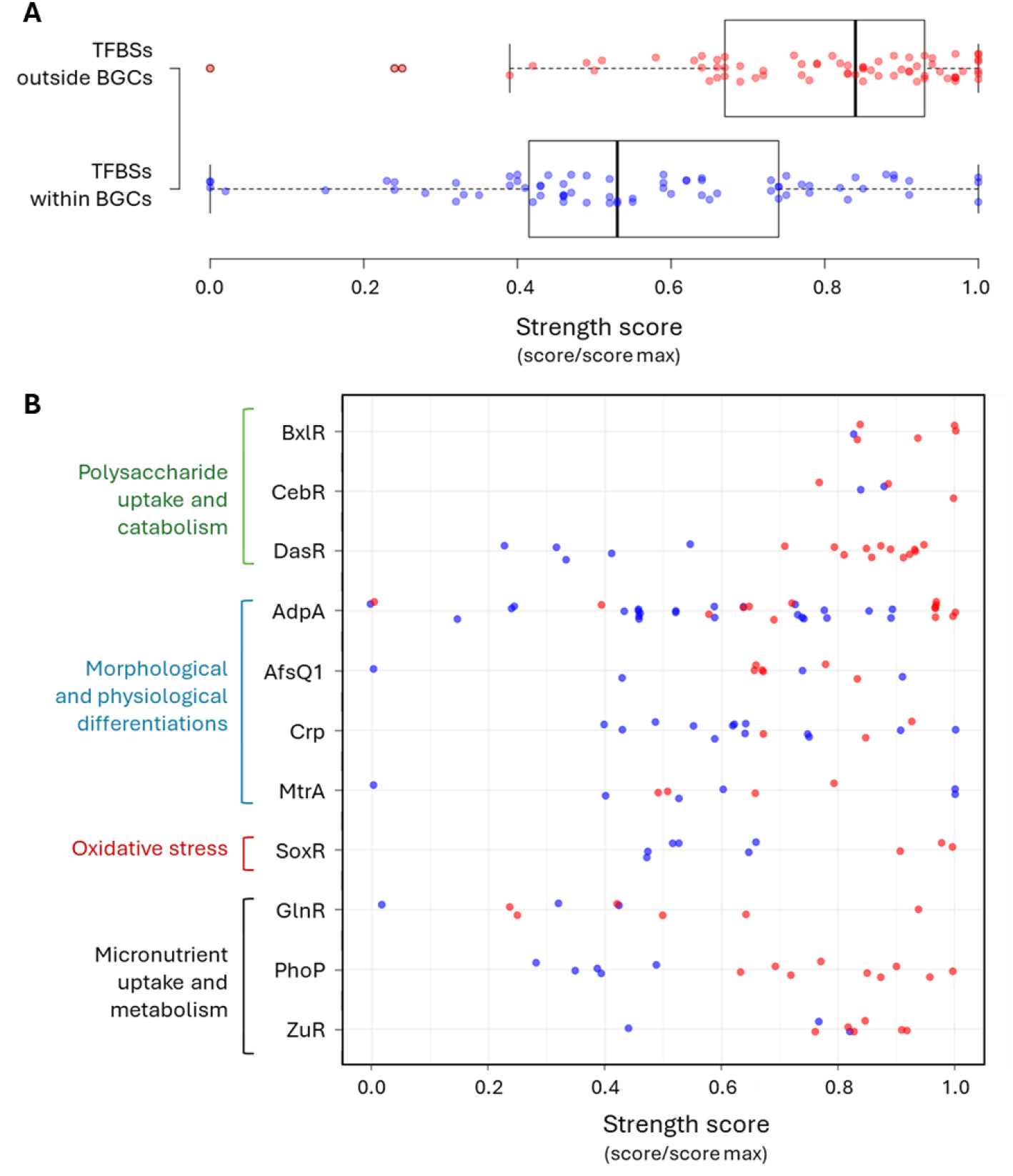
Strength assessment of the TFBS-TF interactions in BGCs and non-BGC genes. **A**. Boxplot showing the interaction strength of TFBSs within BGCs and those associated with non-BGC genes. The edges in the boxplot indicate the 1st and 3rd quartiles (Q), and the median as center line. **B**. Specific distribution of interaction strength scores of 11 global TFs. Blue and red circles are for TFBSs within BGCs and those associated with non-BGC genes, respectively.

## Conclusions

This study sheds light on the complex regulatory architecture governing bacterial BGCs, reflecting a fine-tuned balance between metabolic efficiency, environmental adaptability, and genetic organization. Through a complex interplay of global and pathway-specific transcription factors, bacteria ensure coordinated expression of biosynthetic, transport, and regulatory genes, optimizing resource utilization while maintaining flexibility in response to environmental cues. The observed regulatory patterns of global TFs highlighted a “one-for-all” regulatory strategy by targeting pathway-specific TFs to ensure inclusive transcriptional control over the whole cluster.

Additionally, the variability in transcription factor binding affinities within BGCs suggests a mechanism that allows greater regulatory plasticity, potentially enabling dynamic responses to shifting ecological conditions. A comprehensive understanding of the regulatory networks and the signaling molecules that influence BGC expression is crucial as a strategy for activating pathways leading to novel compound discovery. Bioinformatic tools such as MiniMotif^51^ facilitate the discovery of signaling pathways to BGC expression by cracking the regulatory codes ^21^. To reliably predict how BGCs are controlled, our study shows that parameters other than the similarity of a TFBS to sequences known to be bound by a TF must be considered. The predicted interaction strength is itself a parameter, whose importance varies according to the flexibility of the TF to bind to degenerate sequences. The other criteria that together improve prediction reliability are: i) the functional category of the genes targeted by a TF (Functional targeting), ii) the position of the TFBS for optimal expression control (Positional distribution of TFBSs), and iii) the gene’s organization to assess the number of genes of the BGCs that might be under the control of a single TFBS. These insights not only deepen our understanding of BGC regulation but also provide a foundation for future efforts to optimize culture conditions and manipulate gene expression, paving the way for the discovery and production of novel bioactive compounds.

## Author contributions

Conceptualization: S.R.M., S.R. – Data curation: All authors. – Software: S.R.M. – Formal Analysis: S.R.M. S.R. – Visualization: S.R.M., S.R. – Supervision: S.R. – Writing – original draft: S.R. – Writing – review & editing: All authors.

## Acknowledgments

The work was supported by FNRS aspirant Grant to S.R.M., and an FNRS-PDR T.0195.23 (40013674) Grant to Y.K.

